# The 3′ end of the human norovirus antigenomic sequence is required for its genomic RNA synthesis by RNA-dependent RNA polymerase

**DOI:** 10.1101/2021.05.17.444519

**Authors:** Takashi Shimoike, Tsuyoshi Hayashi, Tomoichiro Oka, Masamichi Muramatsu

**Author notes:** Correspondence: Department of Virology II, National Institute of Infectious Diseases, 4-7-1, Gakuen, Musashi-Murayama, Tokyo, 208–0011, Japan.

## Abstract

Norovirus genome is a single-stranded positive-strand RNA. To reveal the mechanism underlying the initiation of the norovirus genomic RNA synthesis by its RNA-dependent RNA polymerase (RdRp), we used an *in vitro* assay to detect the complementary RNA synthesis activity. Results showed that the purified recombinant RdRp synthesized the complementary positive-sense RNA from the 100 nt template corresponding to the 3′ end region of the viral antisense genome sequence, but that RdRp did not synthesize the antisense genomic RNA from the 100 nt template, corresponding to the 5′ end region of the positive-sense genome sequence. The 31 nt region at the 3′ end of the RNA antisense template was then predicted to form the stem-loop structure. Its deletion resulted in the loss of complementary RNA synthesis by RdRp. The connection of the 31 nt to the 3′ end of the positive-sense RNA template allowed to be recognized by the RdRp. Similarly, an electrophoretic mobility shift assay further revealed that RdRp bound to the antisense RNA specifically, but the 31 nt deletion at the 3′ end lost the binding to RdRp. Therefore, combining this observation with further deletion and mutation analysis, we concluded that the predicted stem-loop structure in the 31 nt and region close to the antisense viral genomic stem sequences are important for initiating the positive-sense human norovirus genomic RNA synthesis by its RdRp.

## Introduction

Norovirus is a single-stranded positive-strand RNA virus that belongs to the Caliciviridae family. It is a major cause of acute gastroenteritis in humans. Its genome is approximately 7.5 knt long, and it has 4 or 5 bases of the 5′untranslated region (UTR) and encodes 3 or 4 open reading frames. ORF1 encodes the viral nonstructural viral proteins (NS1-2, NS3 [NTPase], NS4 [3A-like protein], NS5 [VPg], NS6 [protease], and NS7 [RNA-dependent RNA polymerase; RdRp]), while ORF2 and ORF3 encode the major (VP1) and minor structural protein (VP2), respectively. A 3′ UTR (30–60 nt in length) exists downstream of the ORF3. Also, the 3′terminus of the genome (downstream of the 3′ UTR) is polyadenylated (Fig. 1A) (1).

**Figure 1.**
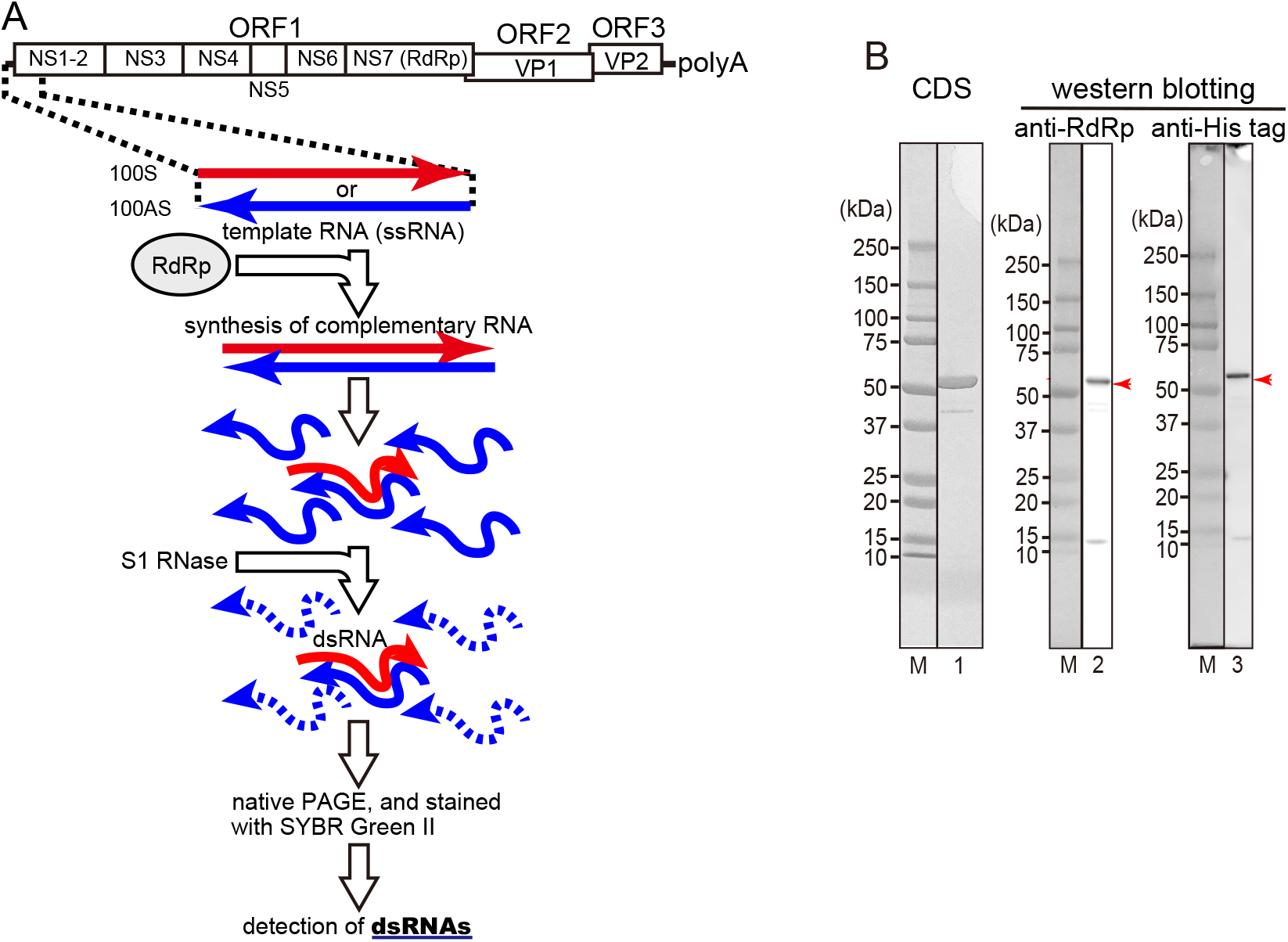
Schematic illustration of a non-radioactive labeling method to detect RNA synthesis by RdRp *in vitro* and confirmation of the purified RdRp. **A:** as shown by red or blue arrows, respectively, the region of the RNA (positive sense or antisense (complementary) sequences of the 5′ or 3′ end 100 nt of the human norovirus genomic RNA) used as templates in this study is shown. Template single-stranded RNA (ssRNA) and norovirus RNA-dependent RNA polymerase (RdRp) were mixed and incubated to synthesize complementary RNA. The newly synthesized ssRNAs formed double-stranded RNA (dsRNA) with an excess of ssRNA templates. The ssRNAs in the reaction mixture were digested with S1 ribonuclease to visualize dsRNA only. RNA molecules before and after S1 ribonuclease treatment were separated by non-denaturing polyacrylamide gel electrophoresis (native PAGE) and stained with SYBR Green II. **B**: confirmation of the purified RdRp. 31.7 nmol (lane 1) or 2.3 pmol (lanes 2 and 3) of the purified RdRp was performed using SDS-PAGE. After SDS-PAGE, the gel was visualized using Coomassie dye staining (CDS; lane 1) or detected with Western blotting using anti-RdRp antiserum (lane 2) or using anti-His-tag antibody (lane 3). M, molecular weight markers.

The viral RdRp is responsible for the replication/synthesis of the norovirus genomic RNA. RdRp first synthesizes antisense RNA using the genomic RNA as a template. It then uses these antisense RNAs to synthesize the progeny viral genomic RNAs. Investigation into the RNA synthesis activity of the norovirus RdRp *in vitro* or in cell-based systems has been reported (2–7). However, the mechanism of replication of the viral genome RNA by its RdRp remains unclear.

In this study, the viral genomic RNA template-specific RNA synthesis by the human norovirus (HuNV) RdRp was demonstrated for the first time in an *in vitro* assay. Results showed how RdRp recognizes the antisense genomic RNA to initiate the synthesis of the complementary positive-sense genomic RNA.

## Results

### HuNV RdRp synthesizes the viral positive-sense RNA with the antisense template RNA

We used a non-radioactive labeling method to detect RNA synthesis by RdRp *in vitro* (a schematic of the procedures is shown in Fig. 1A). After the template RNA was incubated with RdRp, RNA complementary to the template RNA was synthesized, forming a double-stranded RNA (dsRNA) with the template RNA. The dsRNA then remained after incubation with an S1 ribonuclease, to digest the single-stranded RNA into small nucleotides. Subsequently, the synthesis of the dsRNA is then used as an index that RdRp synthesizes the complementary RNA in the following experiments.

In the next steps, the HuNV GII. P3 RdRp tagged with six histidines at its N-terminus was prepared (purity: 96%; lane 1 in Fig. 1B). The 5′ terminal nt 1–100 of viral genomic RNA of the HuNV GII. P3 strain (100S [sense) or its complementary sequence (100AS [antisense) was then used as template RNAs (cf. Fig. 1A). All the template RNAs used here carry GGG and CCC sequences at their 5′ and 3′ ends, respectively, to synthesize template RNAs efficiently *in vitro* and to form the stable dsRNAs. When the increasing amounts of RdRp were incubated with 100S RNA, almost no RNA was synthesized (lanes 4–7 on the upper column in Fig. 2A). However, when the increasing amounts of RdRp were incubated with 100AS RNA, dsRNA was synthesized in increasing amounts (lanes 8–11 in the upper column in Fig. 2A). To confirm that these synthesized RNAs were dsRNAs, the RNAs were treated with S1 ribonucleases. The amounts of the synthesized dsRNA were then increased according to the amounts of RdRp (lanes 8–11 in the lower column in Fig. 2A and the graph in Fig. 2B). The efficient activity of S1 ribonucleases was confirmed by the fact that the marker ssRNAs (100S and 100AS) were digested well, but the dsRNA (ds100) markers were resisted with S1 ribonuclease treatment (lanes 1, 2, and 3 on the lower column in Fig. 2A, respectively). These results therefore indicate that RdRp specifically recognizes the 3′ end of the antisense viral genome RNA and indicate also that this region is important to initiate the synthesis of the positive-strand RNA by RdRp.

### The predicted stem-loop structure and the region close to the stem are required for initiating the synthesis of the positive-sense RNA by RdRp

The 100AS RNA was predicted to form the stem-loop structure at the 3′ terminus (nt 74–87 of the antisense genome RNA of the GII. P3 strain (Fig. 3A). The sequence of the nt 74 to 100 was highly conserved among the norovirus genogroups I, II, IV, and VII (Fig. 3B). To see whether this stem-loop (nt 74–87) was important for initiating RNA synthesis by RdRp, a deletion mutant of 100AS RNA that lacks the 31 nt (nt 70–100) at the 3′ terminus of the 100AS RNA was constructed (designated as Δ31AS RNA, Fig. 4A). Its complementary template RNA was also constructed as a control (designated as Δ31S RNA). When Δ 31AS RNA and Δ 31S RNA were used as template RNAs, RdRp did not synthesize the complementary RNA from either template RNAs, (lanes 11 and 13 at the upper column in Fig. 4A). To remove the single-stranded template RNA to confirm the synthesis of the dsRNA, the reaction products shown on the upper column in Fig. 4A were treated with S1 ribonucleases. Absent or faint RNAs were detected (lanes 11 and 13 in the lower column in Fig. 4A, respectively). These results therefore indicate that nt 70–100 of 100AS RNA containing the predicted stem-loop structure is necessary for initiating the complementary positive-strand genomic RNA synthesis by RdRp.

**Figure 2.**
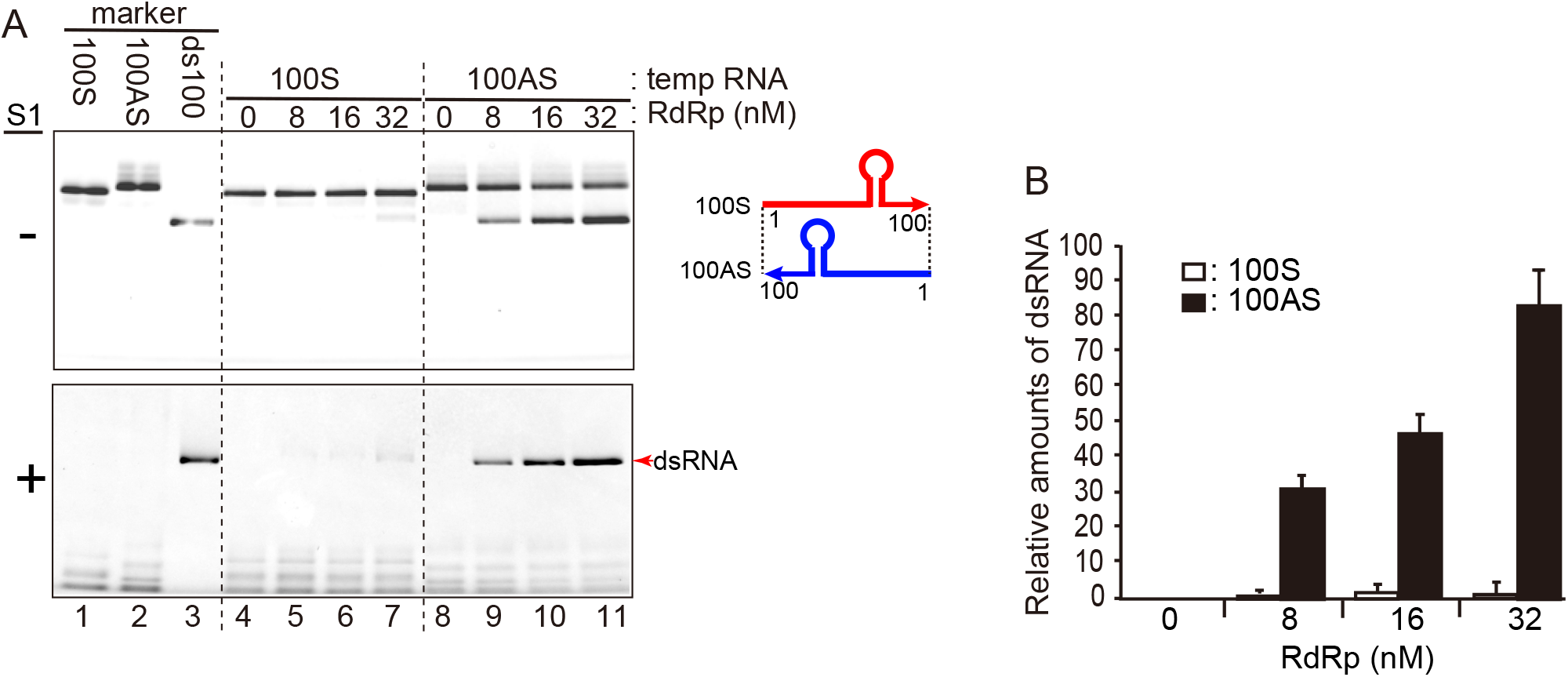
RdRp synthesizes the complementary RNA from 100AS RNA template. **A:***in vitro* synthesized RNAs separated in native PAGE. The template ssRNAs [100S RNA (lane 1), 100AS RNA (lane 2), and ds100 RNA (lane 3) were used as size markers. Five pmol of each template RNA, i.e., 100S (100 nM, lanes 4–7) or 100AS (100 nM, lanes 8–11), and the purified HuNV RdRp (0–32 μM) were mixed. Half of the total volume of samples was mixed with 6× gel-loading buffer and loaded on native PAGE (the upper column). S1 ribonuclease was added to the remaining half of the samples, and the samples were incubated and loaded on native PAGE (the lower column). The dsRNA products separated in native PAGE before and after S1 ribonuclease treatment are shown in the upper and lower columns, respectively. Red arrow indicates newly synthesized RNA products, which had formed dsRNAs with an excess of template RNA and identically migrated to dsRNA markers (lanes 4–11). **B:** the relative amounts of the newly synthesized dsRNA products (the transcriptional activity) were analyzed using ImageJ v. 2.1.0/1.53c computer software and shown as the graph. The amounts shown were the averages of three experiments. The standard deviations are also shown in the graph.

**Figure 3.**
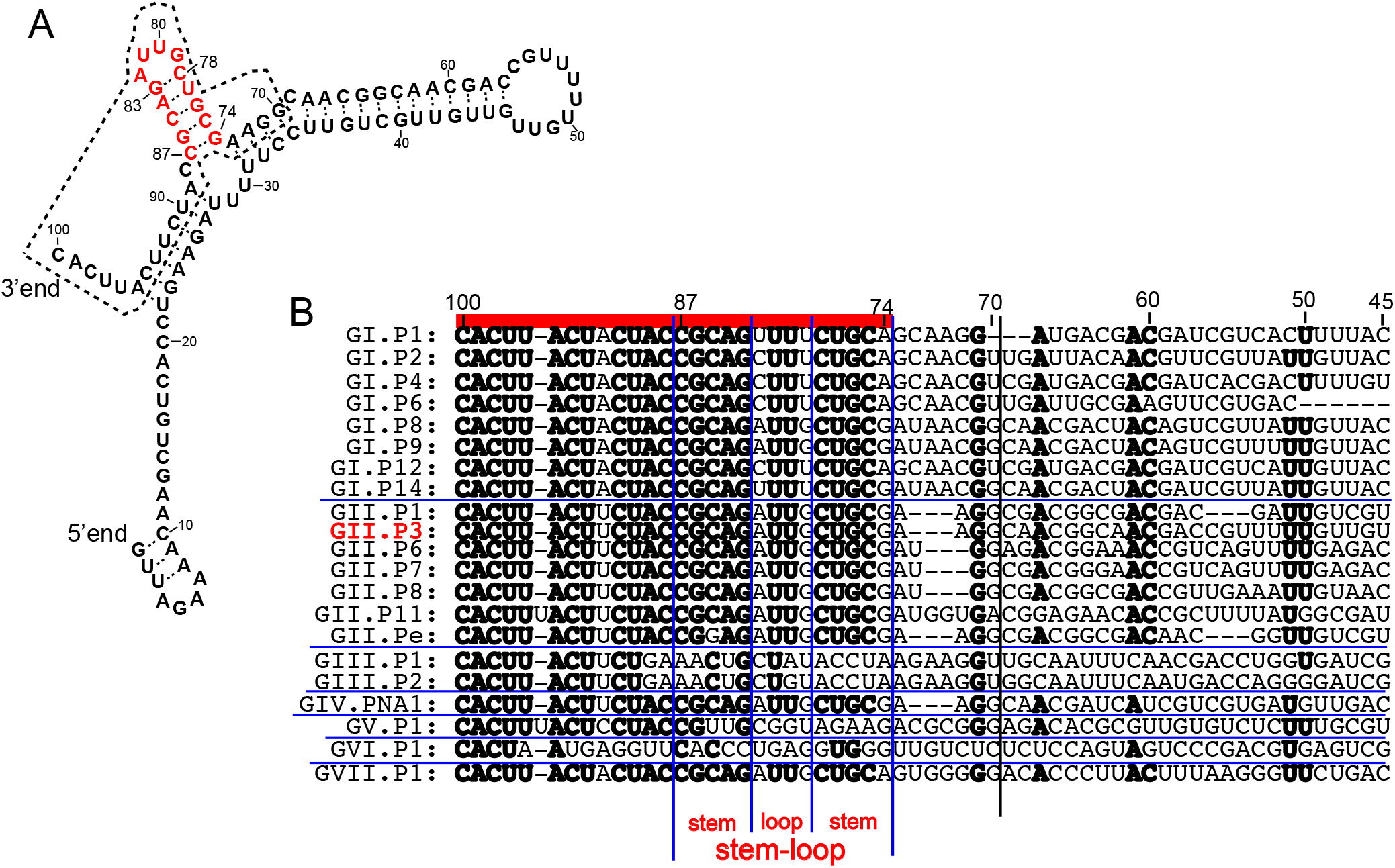
The predicted secondary structure of 100AS HuNV RNA. **A**: the 100AS RNA secondary structure was predicted using the computer software (13). The region of 31AS RNA (nt 70–100 of the U201 strain) is surrounded by a dotted line, and *red characters* indicate nt 74–87, which form the predicted stem-loop structure. B: alignments of 21 kinds of norovirus genotypes at the 3′ end regions of the antisense genome sequences (nt 100–45 of 100AS RNA of the GII. P3 U201 strain and its corresponding regions of other genotypes) are shown. The nucleotides with more than 71% identity (the same nucleotide in more than 15 of the total 21 genotypes) are indicated in *bold*. RdRp-coding genes are used to classify the genotypes (14). *Horizontal blue lines* separate genogroups. R*ed line* on the top indicates highly conserved regions (nt 100–74 regions of the GII.P3 U201 strain). The 31AS RNA region is indicated by separating with a vertical black line between nt 69 and 70. *Four vertical blue lines* show the separation in the stem and loop regions in the stem-loop regions. The GII. P3 U201 strain that was used in this paper is shown in *red characters*.

**Figure 4.**
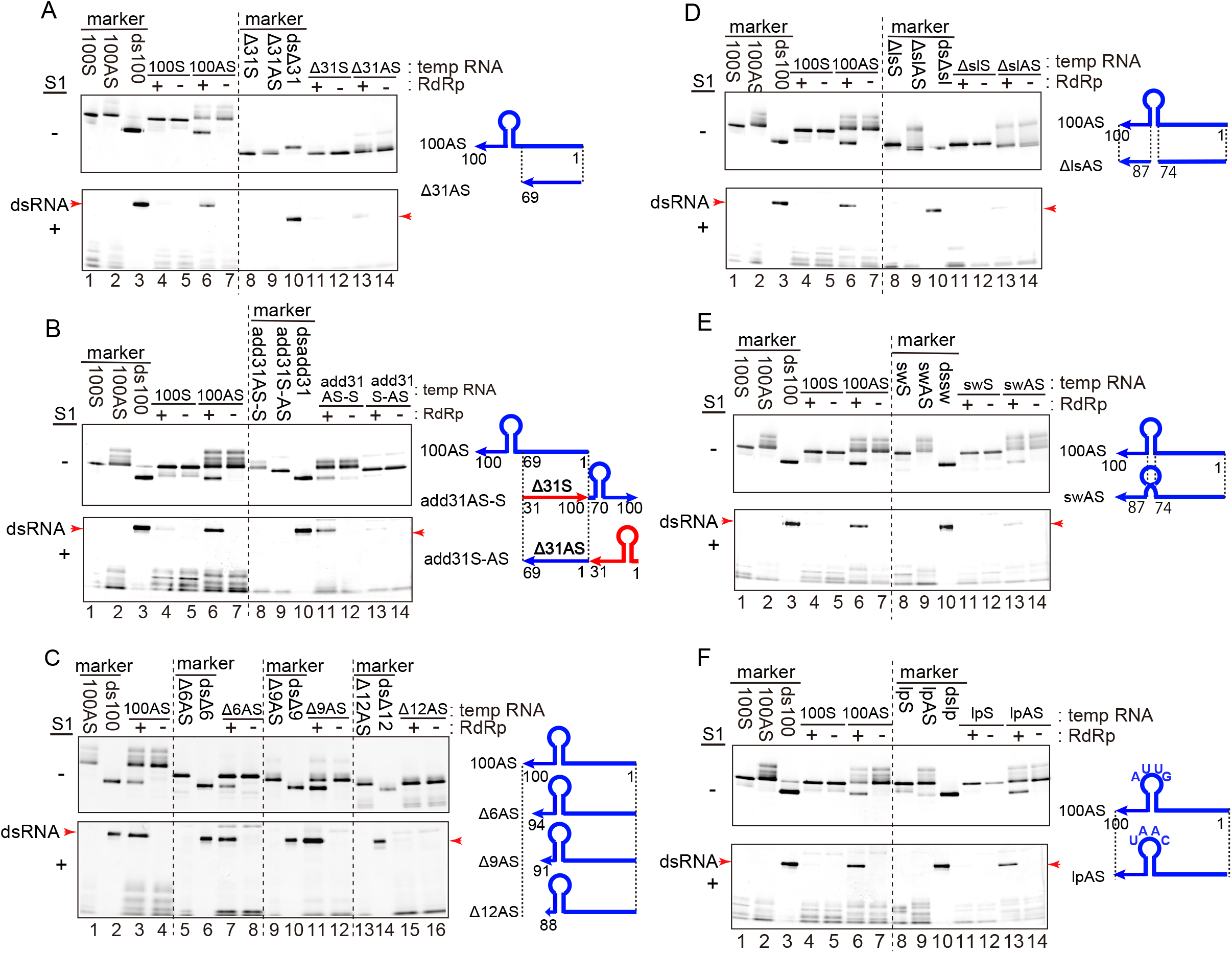
The RNA synthesis with mutated template RNAs by HuNV RdRp. Five pmol of each template RNA (final concentration, 100 nM) and 0.8 pmol HuNV RdRp (final concentration, 16 μM) were mixed. After incubation, half of the samples were loaded on native PAGE (the upper column). The other half of the samples were treated with S1 ribonucleases and loaded on native PAGE (the lower column). The results using templates, Δ 31S and Δ 31AS RNAs (A); add31AS-S and add31S-AS RNAs (B); Δ6AS, Δ9AS, and Δ12AS RNAs (C); ΔslS and ΔslAS RNAs (D); swS and swAS RNAs (E); and lpS and lpAS RNAs (F), were shown, respectively. The template ssRNA (lanes 1, 2, 8, and 9) and dsRNA (lanes 3 and 10) were used as size markers in A, B, and D–F. The template ssRNA (lanes 1, 5, 9, and 13) and dsRNA (lanes 2, 6, 10, and 14) were used as size markers in C. The positions of each dsRNA are indicated with *red arrows* in the lower columns. All results are representative of at least triplicate experiments. The schematic structures of the 100S, 100AS, and its mutated template RNAs are shown on the right side of each figure.

To confirm that the 31 nt RNA, carrying the nt 70–100 of 100AS RNA (31AS RNA, shown in Fig. S1), was sufficient for RNA synthesis by the RdRp, two chimeric RNA templates were further prepared: (i) the 5′ end of 31AS RNA was connected to the 3′ end of the Δ31S RNA (designated as add31AS-S), and (ii) the 3′ end of 31S RNA was connected to the 5′ end of the Δ31AS RNA (designated as add31S-AS and complementary to add31As-S RNA) (cf. Fig. S1). RdRp also synthesized dsRNA with add31AS-S RNA as the template, but a few amounts of dsRNA were detected when the complementary add31S-AS RNA was used as a template (lanes 11 and 13, respectively, in Fig. 4B). These results therefore confirmed the importance of the 31 nt region at the 3′ end of antisense genomic RNA for the RdRp activity.

Similarly, to narrow down the regions in nt 70–100 of the 100AS RNA responsible for initiating RNA synthesis by the RdRp, the following deletion mutants were constructed: Δ6AS, Δ9AS, and Δ12AS RNA, in which nt 95–100, nt 92–100, or nt 89–100 were deleted, respectively (Fig. 4C). The result showed that RdRp synthesized dsRNAs derived from Δ6AS and Δ9AS, respectively, to the same extent as that of the 100AS RNA, whereas the synthesis of dsRNA from the Δ12AS RNA template was much less efficient (Fig. 4C), indicating that the nt 90–100 is not important for initiating dsRNA synthesis.

To know the importance of the stem-loop region (nt 74–87), three kinds of mutant template RNAs were also constructed: (i) Δsl (stem-loop) AS RNA, in which the stem-loop region (nt 74–87) was deleted (Fig. 4D); (ii) sw (<u>sw</u>itch) AS RNA, in which the stem regions, nt 75–78 and nt 83–86, were switched together to preserve the stem structure (Fig. 4E); and (iii) lp (<u>l</u>oo<u>p</u>) AS RNA, in which nt 79–82 (GUUA) at the loop of the stem-loop in the 100AS RNA was changed to the complementary sequence (CAAU) (Fig. 4F). When the ΔslAS and swAS RNAs were used as templates for RdRp, small amount of the dsRNAs was synthesized (lane 13 in Fig. 4D and 4E, respectively). Alternatively, dsRNA was synthesized as well with lpAS RNA (lane 13 in Fig. 4F). These results therefore also indicate that the stem-loop region is important and the stem region (nt 74–78 and nt 83– 87) is more important than the loop region (nt 79–82) (Fig. 3).

### Interaction of the 3′ end of 100 nt antigenomic RNA with RdRp

The direct interaction was examined by the electrophoretic mobility shift assay (EMSA) technique using RNAs labeled with ^32^Phosphate at the 5′ end. When the 100AS RNAs were incubated with RdRp, the amounts of the RNA-RdRp complex increased as the amount of RdRp increased (lanes 1–4 in Fig. 5A). However, when Δ31AS RNA lacking in the 31 nt of the 3′ end of the 100AS RNA was incubated with increasing amounts of RdRp, Δ31AS RNA formed a complex with RdRp much less efficiently than 100AS RNA even at the highest concentration (lanes 5–8 in Fig. 5A). These results indicate that the 31 nt is recognized by RdRp and implies that this recognition causes the initiation of genome positive-strand RNA synthesis.

**Figure 5.**
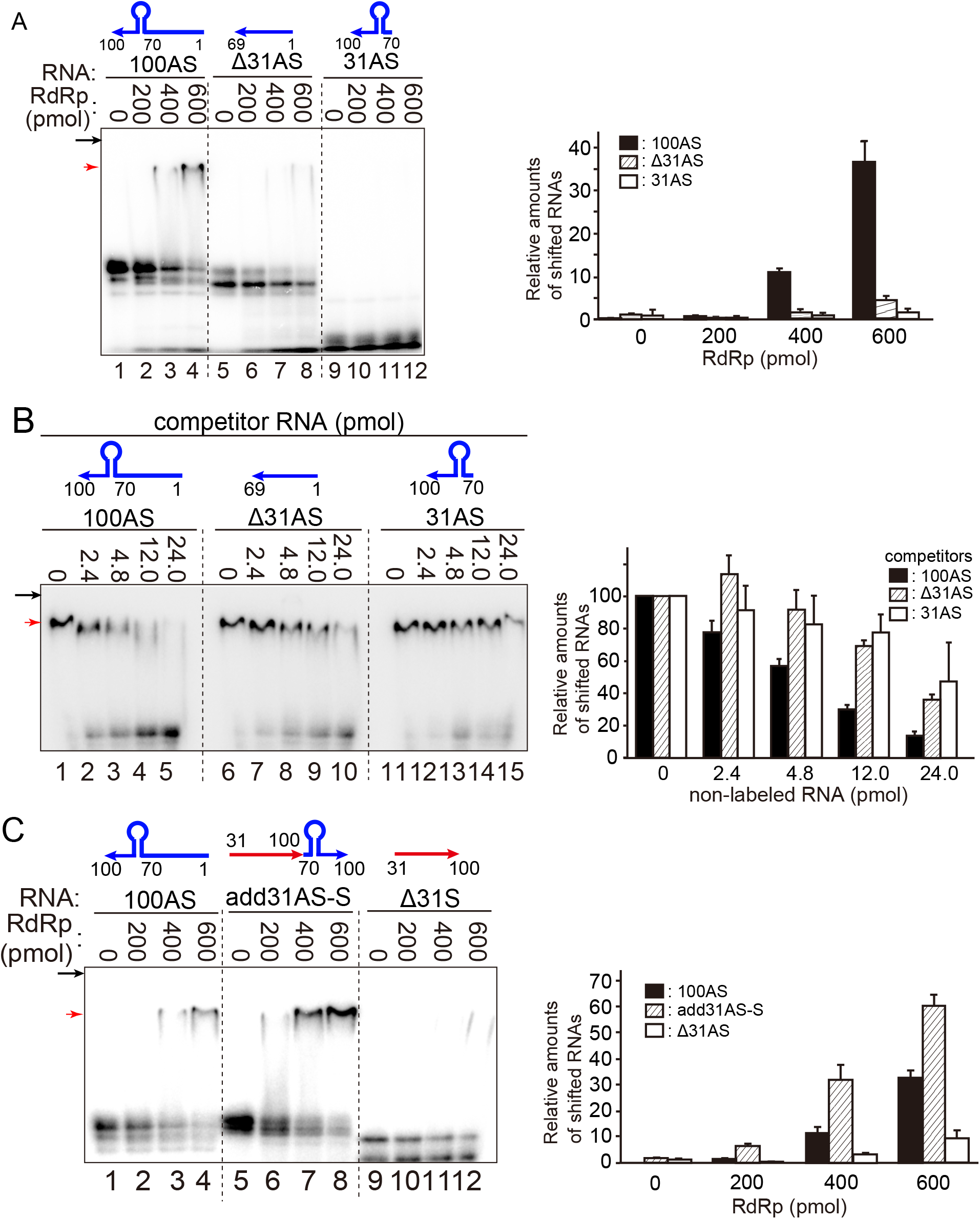
Interaction of the template 100AS RNA and its mutated RNAs with RdRp by EMSA. **A:** the ^32^P-labeled 5′ end-labeled RNAs (100AS; lanes 1–4, Δ31AS; lanes 5–8, or 31AS; lanes 9–12, 1 pmol each) and RdRp (0–600 pmol) were incubated. **B:** the ^32^P-labeled 5′ end-labeled 100AS RNA (0.2 pmol), the indicated number of non-labeled competitor RNAs; 100AS (lanes 1–5), Δ31AS (lanes 6–10), or 31AS RNA (lanes 11–15) and RdRp (600 pmol) were incubated. **C:** the ^32^P-labeled 5′ end-labeled RNAs (1pmol); 100AS (lanes 1–4), add31AS-S (lanes 5–8), or Δ31As (lanes 9–12) RNAs and RdRp (0–600 pmol) were incubated. After incubation, EMSA was used to examine all the samples. The relative number of RNAs that were complexed with RdRp (the shifted RNAs) was analyzed using ImageJ v. 2.1.0/1.53c computer software and shown as the graph. The amounts were the averages of three times experiments. The standard deviations are shown in the graph. Black arrows indicate the position of the bottom of wells. Red arrows indicate the dsRNAs. The schematic structures of labeled RNAs (A and C) or non-labeled competitor RNA (B) are shown on each RNA.

To confirm that this interaction was specific, two non-labeled RNAs, 100AS RNA and Δ31AS RNA, were used as competitors in EMSA (Fig. 5B). When the ^32^Phosphate (^32^P)-labeled 100AS RNA was incubated with increasing amounts of the same non-labeled 100AS RNA, the amount of the labeled RNA-RdRp complex decreased in correspondence with the increase in non-labeled 100AS RNA. The amount of the shifted RNA-RdRp complex decreased to 13.5% (lanes 1–5 in Fig. 5B). In contrast, when ^32^P-labeled 100AS RNA was mixed with increasing amounts of non-labeled Δ31AS RNA lacking in nt 70–100 at the 3′ end of 100AS RNA, the ^32^P-labeled 100AS RNA-RdRp complex competed with non-labeled Δ31AS RNA, but the efficiency of competition with non-labeled 31AS RNA (decreased to 35.8%) was lower than that with the non-labeled 100AS RNA (decreased to 13.5%). These results thus indicate that the 100AS RNA and RdRp have specific interactions and imply that the 31 nt RNA (31AS RNA) is required for this interaction.

Similarly, since the 31AS RNA (the nt 70–100 of 100AS RNA, cf. Fig. S1) was important for initiating genomic RNA synthesis by RdRp as demonstrated above (lanes 13 and 11 in Fig. 4A and 4B, respectively); EMSA was used to examine the interaction of the 31AS RNA with RdRp. The ^32^P-labeled 31AS RNA was incubated with increasing amounts of RdRp. However, no 31AS RNA-RdRp complex was detected even at the highest concentration (600 pmol) of RdRp (lanes 9–12 and the graph in Fig. 5A). Also, when the non-labeled 31AS RNA was used as a competitor for the complex of 100AS RNA and RdRp, the competition was less efficient than when using the non-labeled 100AS RNA as a competitor (decreased to 47.3% *vs*. 13.5%, lanes 11–15 *vs*. lanes 1–5 and the graph in Fig. 5B). It is possible that the 31AS RNA was too short to form the stable complex with RdRp efficiently. Therefore, to determine whether the 31AS RNA region interacted with RdRp, the interaction of RdRp with add31AS-S RNA (the structure formed by connecting the 5′ end of 31AS RNA to the 3′ end of Δ 31S RNA (cf. Fig. S1)) was examined using EMSA. When the ^32^P-labeled add31AS-S RNA was incubated with RdRp, the amount of RNA-RdRp complex increased in correspondence with the amounts of RdRp (lanes 5–8 and the graph in Fig. 5C). However, the Δ31S RNA itself (nt 31–100 of the positive-sense RNA) did not synthesize its dsRNA by RdRp (lane 11 in Fig. 4A). It also did not form a complex with RdRp (lanes 9–12 and the graph in Fig. 5C). Finally, these results show that 31AS RNA is required for interacting with RdRp and required for initiating RNA synthesis as well.

### Inhibitory effects of nucleoside analogs on RNA synthesis by HuNV RdRp

We examined whether our *in vitro* assay detected RdRp inhibitory effects by compounds. We tested 2′-C-methylcytidine (2′ CM) and its ribonucleotide triphosphate form, 2′CM-cytidine triphosphate (2′CM-CTP) as known inhibitors of HuNV RdRp. When increasing concentrations of 2′CM were added to the reaction mixture, no inhibitory effect on RNA synthesis was observed (lanes 3–7 at the lower column and the graph in Fig. 6). When increasing concentrations of the converted form, 2′CM-CTP, were added to the reaction mixture, the amounts of synthesized dsRNA were decreased from 100% to 0.5% (lanes 8–12 and the graph in Fig. 6). Our result was also consistent with previous studies and reasonable because it is known that 2′CM has an inhibitory effect after it is converted to the ribonucleotide triphosphate form, 2′CM-cytidine triphosphate (2′CM-CTP) by cellular enzymes. The IC_50_ (half of the maximal inhibitory concentration) of 2′CM-CTP was calculated as 57 ± 20 μM.

**Figure 6.**
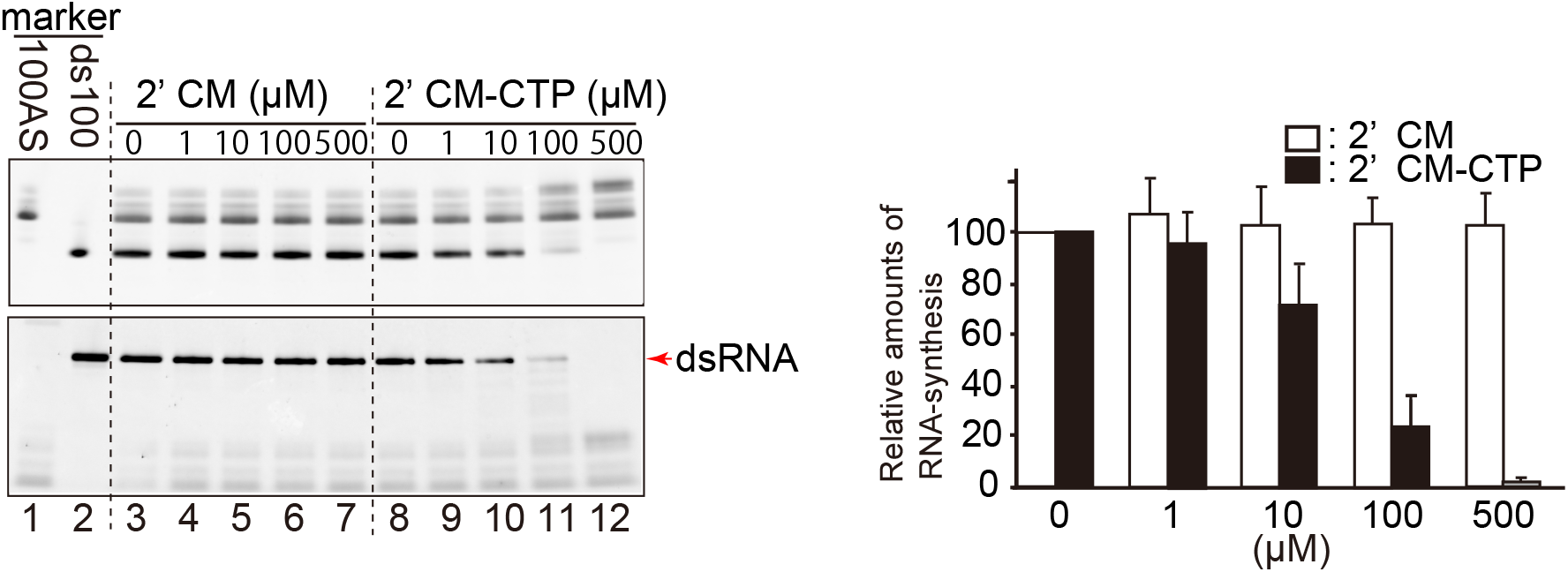
Inhibitory effect of 2′CM-CTP on the RNA synthesis by RdRp. 2′-C-metylcytidine (2′CM; 0–500 μM; lanes 3–7) or 2′C-methylcytidine 5′-triphosphate (2′CM-CTP; 0–500 μM; lanes 8–12) was mixed with 100AS RNA (10 pmol, final concentration: 0.4 μM) and RdRp (45 pmol, final concentration: 1.8 μM). Half of the samples were loaded on native PAGE (the upper column). The other half of the samples were treated with S1 ribonucleases and loaded on native PAGE (the lower column). The template 100AS RNA (lane 1), and ds100 RNA (lane 2) were used as the size markers. Red arrow indicates the dsRNA products. The relative number of dsRNA products is shown in the graph. The amounts were the averages of three times experiments. The standard deviations are also shown in the graph.

## Discussion

From this study, we revealed that the HuNV RdRp initiated the template-specific initiation of RNA synthesis and RdRp interacted with the 3′ end region of the antisense genomic RNA.

First, we demonstrated that HuNV RdRp synthesized complementary (positive-strand genomic) RNA from the 100 nt of the 3′ end region of the antisense genomic template RNA (100AS RNA), but did not synthesize the antisense RNA from the 100 nt of the 5′ end region of the genomic template RNA (100S RNA) (cf. Fig. 2A). Similarly, the introduction of deletion and mutations in the 31 nt region revealed that the predicted stem-loop region and the region close to the stem sequences are required as a template for RNA synthesis by RdRp. There are reports on the activity of HuNV RdRp *in vitro*, and the template RNAs used in these reports are those corresponding to partial 3′ terminal genomic sequences of HuNV (2,3,6), or a non-viral template (4,6,7). HuNV RdRp synthesizes the complementary RNAs by using any sequences of the template RNAs used in these reports. However, there are no specific genomic sequence preferences as template RNAs. In contrast, we demonstrated the genomic sequence specificities of template RNA. The genome specificity would come from the concentrations of RdRp in the reaction mixture; the concentration of RdRp used in these reports was about 30–200 times higher than that used in this study.

Second, we also demonstrated the interaction of HuNV RdRp with the antisense genomic RNA by using EMSA. Results showed that RdRp forms a complex with 100AS RNA and the specific interaction was shown by competition with the same non-labeled 100AS RNA. The deletion of nt 70–100 in 100AS (Δ 31AS RNA) also caused a loss in the interaction with RdRp in EMSA, and the competition with Δ31AS RNA was less efficient than that with 100AS for the 100AS RNA-RdRp binding. These results indicate that the nt 70–100 (31 nt) was required for the interaction with RdRp (Fig. 5A and 5B). The add31AS-S RNA, which is composed of the connection between the 5′ end of 31AS RNA and the 3′ end of Δ31S RNA (shown in Fig. S1), formed the complex with RdRp by EMSA (Fig. 5C). These results also show that the nt 70–100 in 100AS RNA (31AS RNA) is required for interaction with RdRp. As shown in Fig. 4B, the add31AS-S RNA was active as a template for RdRp. Finally, these results imply that the 3′ end (the nt 70–100) of the antisense genomic RNA is required for recognition and initiation by RdRp for the synthesis of the positive-sense genomic RNA.

The nt 74–100 region is highly conserved among genogroups, although the GIII and GVI genogroups are different from this “conserved sequence” (Fig. 3B). This result also indicates that this mechanism of initiation of viral genomic RNA synthesis by RdRp is common among many genogroups, especially the GI, GII, and GIV genogroups, which infect humans.

We also showed that the nucleotide analog, 2′-C-methylcytidine (2′CM-CTP), which was originally found to be one of the inhibitors for the hepatitis C virus (HCV) RdRp, inhibited the synthesis of viral genomic RNA by the RdRp (Fig. 6). The IC_50_ was 57 ± 20 μM. This inhibitory efficiency was similar to that reported by Jin *et al.* (IC_50_ = 33.6 ± 5.8 μM) (6). Similarly, several antiviral drugs have been developed so far, and most of them are direct-acting antivirals (DAAs) that selectively target viral components (e.g., RdRp, helicase, or protease) without affecting cellular function, thus minimizing side effects (8,9). Our *in vitro* RdRp assay system will therefore be a valuable tool to screen seed compounds for developing DAAs against HuNV RdRp.

Also in this study, we identified the regions in the antigenomic HuNV RNA that are required for the interaction and initiation of the genomic RNA synthesis by its RdRp *in vitro*.

### Experimental procedures

#### Plasmid architecture

The plasmid for expressing the RdRp protein coding for the cDNA (nt 3581–5113) of the human norovirus GII. P3_GII.3 U201 strain (accession no. AB0397282) (10) was amplified by PCR using the specific primers containing the NdeI restriction sites at the 5′ end, the stop codon (TAA), and the XbaI restriction site at the 3′ end. Thereafter, it was cloned into the NdeI-XhoI site of the pET28b vector (Merck Millipore, Billerica, MA, USA) (designated pET28b-RdRp).

All plasmids for the template RNAs were prepared by PCR using nt 1– 100 cDNA corresponding to the 5′ terminal of the genomic RNA of the human norovirus GII.P3_GII.3 U201 strain (corresponding to the positive strand) or its complementary sequence (corresponding to the antisense strand), specific primers containing the XbaI restriction site-T7 promoter sequences-three Gs (GGG) at the 5′ end, or three Cs (CCC) at the BsaI-EcoRI sites at the 3′ end (shown in the table at Supporting information), and Ex Taq DNA polymerase (Takara, Shiga, Japan). The amplified DNA fragments were then cloned into the same restriction sites of pUC19 vector (Takara, Shiga, Japan). When the plasmids, pUCadd31AS-S and pUCadd31S-AS, were prepared, pUCΔ31S or pUCΔ31AS was used as the DNA template for PCR. Similarly, all the plasmids for the template RNA were designed to carry GGG and CCC sequences at the 5′ and 3′ ends, respectively. This plasmid was then used to synthesize template RNAs efficiently by T7 polymerase and to form the stable dsRNAs.

#### Preparation of the HuNV RNA-dependent RNA polymerase (RdRp)

*Escherichia coli* cells, ArcticExpress (DE3) (Agilent Technologies, Inc., Santa, Clara, CA, USA) carrying the pET28b-RdRp were cultured in 1 L of Luria Broth Base medium (Invitrogen, Carlsbad, CA, USA) at 30°C until the turbidity at the optical density (OD)_590_ was approximately 0.6 (for up to 3 h) and then placed in ice-cold water for 20 min. Next, isopropyl *β*-D-1-thiogalactopyranoside (final concentration, 1 mM; Wako, Osaka, Japan) was added and the cells were cultured for an additional 24 h at 13°C. To purify RdRp (His-RdRp) containing a His tag at the N-terminus, the culture was then centrifuged at 4400 ×g on a JLA-10.500 rotor (Beckman Coulter, Brea, CA, USA) for 5 min. Afterward, 20 ml of buffer (50-mM Tris-HCl (pH 7.5), 150-mM NaCl, and 5-mM β-mercaptoethanol) was added to the bacterial pellets and the pellets in the buffer were sonicated by five rounds of a 30 s on ¾-in. chip at an intensity level of 5 with subsequent cooling in iced water (Astrason Ultrasonic Processor XL-2020; Misonix, Inc., Farmingdale, NY, USA). The sample was then centrifuged at 22700 ×g on a JA-20 rotor (Beckman Coulter) for 15 min, after which the supernatants were loaded onto a 2.5-mL buffer-equilibrated Ni-NTA column (Qiagen, Hilden, Germany). After washing the column first with 200-mL buffer, and then with 15-mL His-A buffer (50-mM Tris-HCl (pH 7.5), 150-mM NaCl, 5-mM β-mercaptoethanol, 10-mM imidazole), His-RdRp was eluted with 10-mL elution buffer (50-mM Tris-HCl (pH 7.5), 150-mM NaCl, 5-mM β-mercaptoethanol, 200-mM imidazole). The His-RdRp was subsequently purified further with a buffer-equilibrated gel filtration column (HiLoad 16/60 Superdex 200; GE Healthcare, Little Chalfont, UK).

#### Confirmation of RdRp

Here, 31.7 nmol (lane 1) or 2.3 pmol (lanes 2 and 3) of the purified RdRp was analyzed by SDS-PAGE. After SDS-PAGE, lane 1, the gel stained with Coomassie dye, GelCode Blue Stein Reagent (ThermoFisher Scientific, MO, USA) was used to visualize the purified RdRp. Similarly, lanes 2 and 3, proteins on the gel were transferred to the polyvinylidene fluoride (PVDF) membrane with iBlot Gel Transfer Stacks PVDF, regular (#IB401001, ThermoFisher Scientific, MO, USA). The two PVDF membranes were then blocked with PVDF-blocking reagent for CanGet Signal (TOYOBO, Tokyo, Japan) for 1 h at room temperature (RT). One PVDF membrane (lane 2) was incubated with 1/1000 diluted anti-RdRp antiserum (anti-Guinea Pig, polyclonal against RdRp of GII.P3U201 strain, Eve bioscience, Wakayama, Japan). The other PVDF membrane (lane 3) was incubated with 1/1000 diluted anti-6X His-tag antibody (anti-6X His-tag antibody #ab9108; Abcam, Cambridge, MA) in a Can Get Signal Immunoreaction Enhancer Solution 1 (#NKB − 201; TOYOBO, Tokyo, Japan) at 37 °C for 1 h, respectively. The two PVDF membranes were washed with TBS Tween20 (10-mM Tris-HCl [pH 7.5, 150-mM NaCl, 0.05% Tween20) for 10 min three times, after which the PVDF membrane incubated with anti-RdRp serum, or the other PVDF membrane incubated with anti-His tag antibody were then incubated with antibodies: 1/30000 diluted Peroxidase-AffiniPure Donkey Anti-Guinea Pig IgG (H + L) (#706-035-148; Jackson ImmunoResearch Laboratories, Philadelphia, PA, USA), or 1/30000 diluted Goat anti-Rabbit IgG (H+L) HRP-conjugated (#1706515; Bio Rad, Hercules, CA, USA) in a Can Get Signal Immunoreaction Enhancer Solution 2 (#NKB − 301; TOYOBO, Tokyo, Japan) at RT for 1 h, respectively. After this step, the two PVDF membranes were washed with TBS Tween20 for 10 min three times again and then detected with a Super Signal WestFemto Maximum Sensitivity Substrate (#34096; Thermo Fisher Scientific, MO, USA). The RdRp was detected using the LAS-3000 imaging analyzer (Fujifilm, Tokyo, Japan). The purity of RdRp was subsequently calculated using ImageJ v.2.1.0.

#### Preparation of template RNAs

Fifty micrograms of each plasmid were digested using BsaI (New England Biolabs, Ipswich, MA, USA) to exclude the non-viral sequences at the 3′ end. RNA was then synthesized from a template of the linearized DNA using T7 polymerase and purified with a gel filtration column (HiLoad 16/60 Superdex 75, or 200) (the purity of RNAs; at least 90%) as described by cf. Mckenna *et al.* (11) and Shimoike *et al.* (12).

The 5′ end ^32^P-labeled template RNAs were prepared as follows: the calf intestinal alkaline phosphatase (Takara, Shiga, Japan) was used to dephosphorylate the 5′ end of the purified template RNAs. Subsequently, the ^32^Phosphate was bound to the 5′ end of the dephosphorylated template RNA by the T4 polynucleotide kinase (Takara, Shiga, Japan), followed by phenol/chloroform, and precipitated by ethanol. The unreacted γ-32P-ATP in the RNA was then removed using Sephadex G-50 fine resin (GE Healthcare, WI, USA). The ^32^P-labeling efficiencies of RNAs were found to be 49%–51%.

#### Assay for RNA synthesis by RdRp

A 5-pmol template RNA (final concentration, 100 nM) and 0.8-pmol HuNV RdRp (final concentration, 16 nM) were mixed with the reaction buffer (20-mM Tris-HCl [pH 7.4, 5-mM DL-Dithiothreitol, 2-mM NTPs (ATP, UTP, GTP, CTP; Sigma Aldrich, St. Louis, MO, USA)), 2-mM MnCl_2_, 0.8-U/μL ribonuclease inhibitor (Toyobo, Osaka, Japan)) and incubated at 30°C. The mixture was divided into two portions (20 μl each) 30 min after incubation. To one portion, 4 μL of 6× gel-loading buffer (10% glycerol, 0.02% BPB, 0.01% xylene cyanol in TBE buffer [89-mM Tris, 89-mM borate, 2-mM EDTA) was added, and non-denaturing polyacrylamide gel electrophoresis (native PAGE) with the running buffer (25-mM Tris, 192-mM glycine, pH 8.3) was performed at a constant voltage of 300 V for 25 min (5%–20% polyacrylamide gel [ePAGEL, ATTO, Tokyo, Japan). To the other portion, 7 μL of 10-times diluted S1 nuclease (89 U/μL; Promega, Madison, WI, USA) and 3 μL of S1 ribonuclease buffer was added and incubated for 30 min at 30°C to degrade the ssRNA and detect double-stranded RNA (dsRNA) comprising the template RNA and the newly synthesized complementary strands. After this step, the 6× gel-loading buffer was then added, and the native PAGE was performed as described above. After native PAGE, the gel was then stained with SYBR Green II (LONZA, Basel, Switzerland) and detected using LAS-3000 imaging analyzer (Fuji Film, Tokyo, Japan).

#### Prediction of the RNA secondary structure and alignments of NV antisense genome sequences

A. The nucleotide secondary structure of the 100AS RNA (nt 1–100, GII.P3 U201 strain) was predicted using the Genetyx-Mac software v.19.0.3 (Genetyx Corporation) by “RNA secondary structure prediction” (13) using default settings. B. The alignments of the 3′ end 100 nt regions of the norovirus antisense genomic RNA sequences were then calculated using the Genetyx-Mac software v.19.0.3 (Genetyx Corporation) from “multiple alignments” with default settings. The nt 100–45 regions are shown. Similarly, the accession numbers of these 21 sequences are as follows: GI. P1: M87661; GI. P2: L07418; GI. P4: AB042808; GI. P6: AF093797; GI. P8: KJ196298; GI. P9: KF586507; GI. P12: AB039774; GI. 14: AB187514; GII. P1: U07611; GII. P3: AB0397282 (used in this paper); GII. P6: AB039778; GII. P7: AB039777; GII. P8: AB039780; GII. P11: AB126320; GII. Pe: AB434770; GIII. P1: AJ011099; GIII. P2: AF097917; GIV. PNA1: KX907728; GV. P1: FJ875027 (MNV-1); GVI. P1: FJ875027; and GVII. P1: FJ692500.

#### Electrophoretic mobility shift assay (EMSA)

The ^32^P-labeled 5′ end-labeled RNA (1 pmol) and RdRp (200, 400, 600 pmol) were incubated in the reaction buffer (50-mM Tris-HCl pH 7.4, 1.7% glycerol, and 5-mM β-mercaptoethanol) at 30°C for 10 min (total volume, 20 μL). A 4 μL of 6× gel-loading buffer was then added to the reaction mixture. The RNA and RdRp complex was separated with native PAGE at a constant voltage of 60 V for 90 min at 4°C in the 1/2× running buffer (12.5-mM Tris, 86-mM glycine, pH 8.3) after the gel was pre-run at a constant voltage of 60 V for 30 min at 4°C and analyzed by Typhoon FLA 7000 image analyzer (GE Healthcare, WI, USA). In the case of the competition assay, the procedures are the same as above except that the non-labeled RNA (0, 2.4, 4.8, 12.0, or 24.0 pmol) was incubated with ^32^P-labeled 100AS RNA (0.2 pmol) and RdRp (600 pmol), respectively.

#### Assay for inhibitory effects of the nucleoside analog on RNA synthesis

2′-C-methylcytidine (2′ CM, Merck, Germany) and 2′-C-methylcytidine triphosphate triethylammonium salts (2′CM-CTP, Carbosynth Limited, UK) were added (final concentration of 0, 1, 10, 100, or 500 μM) to the reaction mixture that is the same content described in the *Assay for RNA synthesis by RdRp* and incubated at 30°C for 60 min, after which the dsRNA signal was detected as described above. The chemical compound concentrations resulting in a 50% reduction in dsRNA drug-free control production were determined based on the dose-response curve and defined as the mean 50% inhibitory concentration (IC_50_) values of the RdRp activity.

## Data availability

All data are contained within the manuscript

## Supporting information

This article contains supporting information

## Acknowledgment

We are grateful to Dr. Yoshiaki Okada and all the members of our department for their assistance. We also thank Dr. Kazuhiko Katayama for the RNA of the norovirus GII.P3_GII.3 U201 strain.

## Funding and additional information

This study was supported in part by grants from the Ministry of Health, Labour, and Welfare of Japan and the Research Program on Emerging and Re-emerging Infectious Diseases program from the Japan Agency for Medical Research and Development (JP16fk0108304) to T.S.

## Abbreviations

HuNV: Human Norovirus
RdRp: RNA-dependent RNA polymerase
UTR: untranslated region
NS: non-structural protein
S: sense
AS: antisense
EMSA: electrophoretic mobility shift assay
2’CM: 2′ -C-methylcytidine
2’CM-CTP: 2′ CM-cytidine triphosphate
IC_50_: half of the maximal inhibitory concentration
PVDF: polyvinylidene fluoride

